# Virion-wide interactome mapping of HSV-1 reveals maturation-dependent remodeling and convergent organization of herpesvirus tegument networks

**DOI:** 10.64898/2026.08.07.743351

**Authors:** Lars Mühlberg, Yannick Jensen, Julia Ruta, Iris Gruska, Jens B. Bosse, Lüder Wiebusch, Fan Liu, Boris Bogdanow

## Abstract

Herpesvirus virions form by remodeling of intracellular virus-host interaction networks into evolutionarily conserved particle architectures. Here, we define a virion-wide spatial and quantitative protein proximity map of herpes simplex virus 1 (HSV-1) by combining cross-linking mass spectrometry with quantitative proteomics. Integration with intracellular interaction maps reveals that maturation acts as a selective filter, transforming broad virus-host associations into an organized virion network. This process depletes biosynthetic and nuclear components while enriching interactions involved in tegument organization and envelope acquisition around the viral protein UL49. Comparison with analogous maps of human cytomegalovirus (HCMV) identifies HSV-1-UL49 and HCMV-UL32 as functionally equivalent network hubs despite lacking evolutionary relatedness. Both hubs converge on shared phosphoregulatory host factors, short linear interaction motifs, and liquid-liquid phase separation. At the virion surface, the host complement regulator CD59 protects particles from complement-mediated inactivation. Together, these findings show how conserved organizational principles shape virus-specific virion interaction networks during herpesvirus maturation.

## Introduction

The de novo assembly of infectious virus particles is a central step in the viral replication cycle. Viral proteins are often multifunctional, engage in extensive interaction networks, and are produced in quantities that far exceed their final abundance in mature virions. How this complex intracellular interaction landscape is reorganized and condensed into mature virions with defined architecture and stoichiometry remains a fundamental question in virology.

Addressing this question is particularly challenging in herpesviruses. These large double-stranded DNA viruses assemble structurally complex, enveloped particles through a tightly coordinated, multi-step intracellular process (*1*). Assembly begins in the nucleus with the formation of an icosahedral capsid (*2*) and subsequent packaging of the viral genome through a dedicated capsid portal. Following nuclear egress, capsids transit to the cytoplasm, where they undergo tegumentation and envelopment, acquiring a dense, protein-rich tegument layer (*3, 4*) and a Golgi-derived membrane envelope enriched in viral glycoproteins (*5–7*).

Despite substantial genetic diversity, herpesviruses preserve a common virion morphology through a combination of canonical core and more variable, lineage-specific genes (*8*). Core genes predominantly encode protein components required for DNA replication, genome packaging, capsid assembly and nuclear egress. In contrast, tegument and envelope proteins are generally less conserved and often contribute specific functions (*9, 10*), including modulation of host pathways and selective recruitment of host factors into virions (*3*). How evolutionary divergent herpesviruses converge on a structurally and functionally coherent tegument layer, capable of coordinating capsid trafficking, envelopment, and early infection events, remains incompletely understood.

Here, we apply proteome-wide cross-linking mass spectrometry (XL-MS) to intact extracellular particles of the human alphaherpesvirus herpes simplex virus type-1 (HSV-1), in order to construct a proximity map that resolves physical protein-protein contacts across all virion layers. Integrated with quantitative proteomics, this framework defines the composition and spatial organization of viral and host components within the mature virion and identifies the tegument protein UL49 (VP22) as a central interaction hub. Comparison with a similar proximity map of the human betaherpesvirus cytomegalovirus (HCMV) (*11*) reveals convergent principles of tegument organization. Specifically, the non-orthologous proteins HSV-1-UL49 and HCMV-UL32 (pp150) occupy analogous hub positions, reflecting shared interaction partners and biophysical properties. By integrating the virion map with intracellular HSV-1 interaction landscapes, we further delineate how broader protein interaction networks are selectively filtered and remodeled during virus maturation (*12*). Finally, we identify CD59 as a host factor that is selectively recruited to the outer surface of the virion envelope and protects HSV-1 particles from complement-mediated lysis. Together, these findings provide a molecular, network-based view for understanding how conserved and virus-specific interactions shape herpesvirus assembly and maturation.

## Results

### Building the HSV-1 virion interactome

To gain insight into the interactome of HSV-1, we applied virion-XL-MS (*11*) to intact extracellular particles of HSV-1 (**Figure 1A**). Two biological replicates showed high overlap in detected PPIs (**Supplementary Figure 1A**), giving rise to the final dataset of 239 PPIs that have been detected in both replicates (**Figure 1B, Supplementary Table 1**). Of those, 34 (∼14.2 %) had direct prior evidence in the literature in focused studies (**Supplementary Table 2**). For example, we captured the known virion-host associations of eukaryotic initiation factor 4 H (EIF4H) with tegument protein UL41 (Vhs) (*13*) and the complex of UL46 (VP11/12) and host Growth factor receptor-bound protein 2 (Grb2) (*14*).

**Figure 1:**
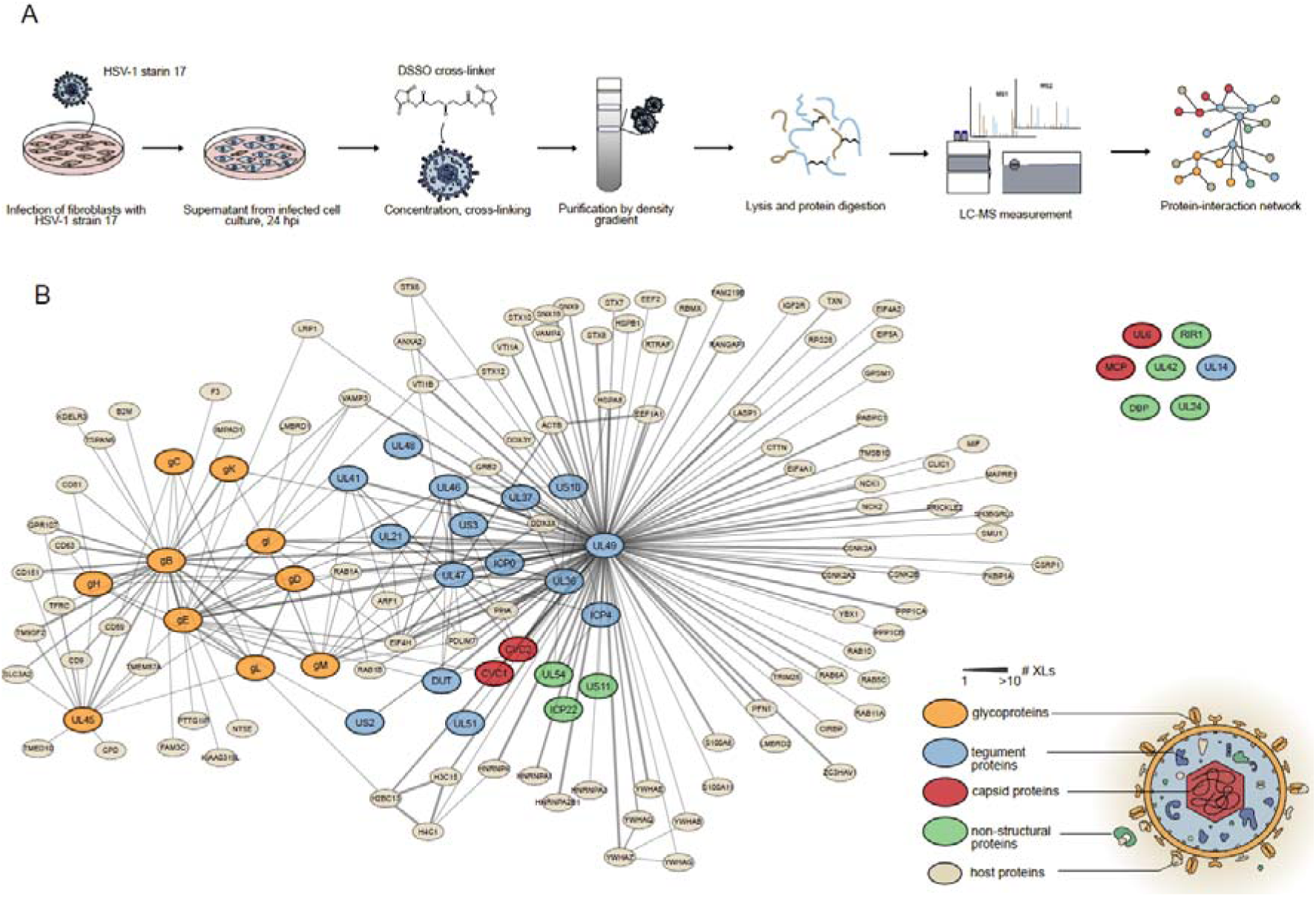
Building the virion-wide HSV-1 protein interactome. **A:** Schematic representation of the workflow used to derive a virion-wide PPI map of Herpes simplex virus type 1. Virions are harvested from infected fibroblast cell culture supernatant at 24 hours post infection (hpi). Virions were concentrated by centrifugation and cross-linked using disuccinimidyl sulfoxide (DSSO). **B:** HSV-1 virion protein interaction network derived from the overlap of two biological replicates following the XL-MS workflow. Nodes represent individual proteins and edges represent protein associations detected by one or multiple cross-links (line thickness). Nodes are color-coded according to protein classes: orange: viral glycoproteins, blue: viral tegument proteins, red: viral capsid proteins, green: viral non-structural proteins, beige: host proteins. Intra-links (cross-links within the same protein) are removed for better visibility.

We then asked whether the identified cross-links faithfully reflect the known architecture of HSV-1. As expected for DSSO (*15*), cross-links of transmembrane proteins remained leaflet-specific and only a minimal amount (1.7 %, **Supplementary Figure 1B, Supplementary Table 3**) connected protein domains from opposing sides. All apparent topology violations originate from glycoprotein E (gE). While the molecular basis of these gE-associated exceptions remains unclear, their restriction to one glycoprotein suggests a protein-specific phenomenon rather than a global artifact of virion cross-linking. Additionally, when mapped on high-resolution Cryo-EM structures of trimeric glycoproteins B in pre-fusion (pdb 9Q9L (*16*)) and post-fusion (pdb 5V2S (*17*)) conformation, approximately 83 % of links were in agreement with the maximum C_α_-C_α_ cross-linking distance of 40 Å **(Supplementary Figure 1C, Supplementary Table 3)**. This suggests that both pre- and postfusion conformation of gB exist on virions, as also shown for HCMV (*18*).

Finally, we assessed whether the cross-links agree with the known layered architecture of HSV-1 virions. Consistent with the known architecture of HSV-1, tegument proteins were connected to all layers, while glycoproteins and capsid shared only few links. Non-structural proteins were exclusively found with tegument proteins, while host proteins cross-linked to all layers **(Supplementary Figure 1D, Supplementary Table 3)**. Taken together, our cross-linking workflow derived reproducible high-confident PPI data that agree with the known topology and protein structures of HSV-1 virions.

### Quantifying selectively recruited host proteins

The network contains several proteins that fulfill their role in the infected cell, e.g. UL54, UL30/UL42 DNA polymerase (non-structural proteins). We hypothesized that these non-structural proteins may only arrive in the particle due to their high intracellular abundance. By contrast, biologically relevant components of the virion should be selectively recruited and thus be enriched in the virion relative to the infected cell.

Aiming to discriminate between selectively recruited proteins and by-standers, we compared protein levels in purified virus particles to infected cells by label-free quantitative proteomics. Indeed, we found that all viral structural proteins had high enrichment ratios, while non-structural proteins showed a lower relative abundance in the virion than in the cell (**Supplementary Figure 2A**). Interestingly, host proteins displayed diverse fold-changes. For example, EIF4H was over 64-fold enriched over the infected cell, while actin, heat-shock proteins or histones were not enriched or even depleted from the particles (**Supplementary Table 4**).

We additionally determined absolute per-virion-copy numbers of all incorporated proteins through regression analysis using the known copies of capsid proteins derived from high-resolution structures (*19*) as an internal standard (**Supplementary Figure 2B, Supplementary Table 4**). This shows that an HSV-1 particle contains ∼ 12,500 protein copies on average, of which ∼3,000 are host-derived. 121 out of the 141 proteins present in our PPI network have average copy numbers of above one, which means they can be considered constitutive components of the virion. About 65 % of the virion proteome was captured in both XL-MS replicates, and 75% was represented by cross-links in at least one replicate (**Supplementary Figure 2B, Supplementary Table 4**). The largest proportion of protein copies absent from our PPI data can be attributed to small capsomer protein (SCP, ∼1300 copies), triplex capsid protein 1 and 2 (TRX2, ∼580 copies and TRX1 ∼220 copies) and cytoplasmic envelopment protein 2 (UL16, ∼180 copies). These viral proteins contain no or only few lysine residues and are therefore hardly detectable using our workflow.

Comparing enrichment levels to copy numbers, we found that 30 viral and 37 host proteins in the PPI-map were present with fold-changes > 2 and copy number > 1 (**Figure 2, Supplementary Figure 2C**). On average, host proteins that directly cross-linked to viral proteins enriched 10-fold stronger with viral particles than other host proteins (**Supplementary Figure 2D**). This provides evidence that host protein enrichment is associated with specific interactions with viral proteins and suggests that the combined information of protein proximity and enrichment allows recapitulating recruitment mechanisms.

**Figure 2:**
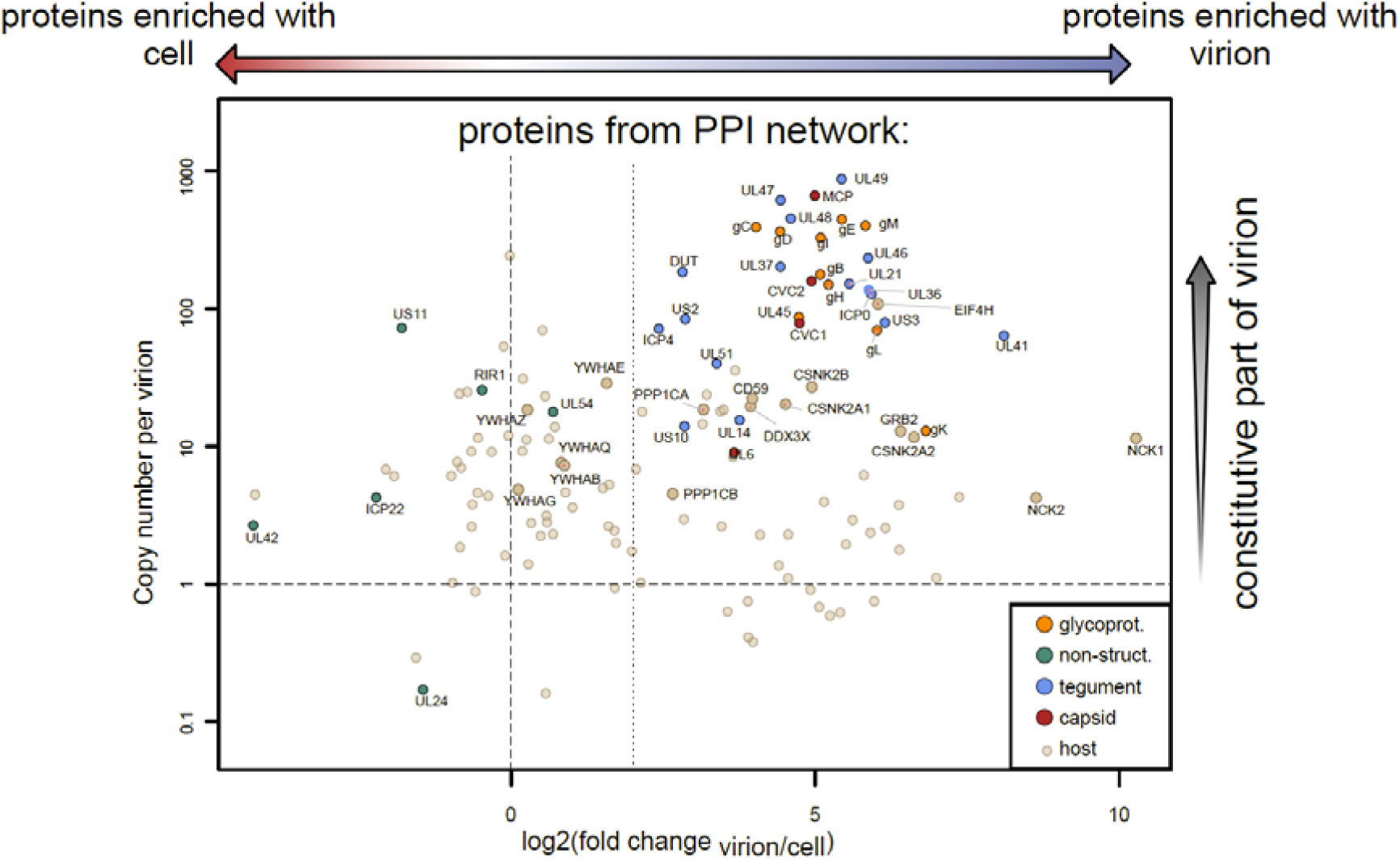
Quantifying the cross-linked proteome of HSV-1 virions. Scatterplot shows relative enrichment ratios (log2 fold change from LFQ values, virion over infected cell) over the absolute per-virion copy number (on a decimal logarithmic scale) of proteins from the PPI-network. Points are color-coded according to Figure 1B. Cut-offs for protein abundance (1 protein copy per virion) and for enrichment (same relative abundance and 4-fold enrichment in the virion) are indicated by dashed lines. All viral proteins as well as selected host proteins are labeled by gene name. For a fully annotated representation see Supplementary Figure 2C.

### Divergent herpesviruses converge on analogous tegument hub organization

The recruitment of host proteins into virions is a common characteristic of Herpesviruses (*20*). We therefore asked whether the interactions with host proteins are conserved between the alphaherpesvirus HSV-1 and the beta-herpesvirus HCMV, which we studied earlier using similar proteomics workflows and the same type of host cells (*11*). Overall, we found that host proteins contributed a larger share of protein copies in HSV-1 than HCMV. Nevertheless, host-protein enrichment ratios were moderately correlated between the two viruses (Spearman’s ρ ≈ 0.59), indicating that many host proteins are incorporated into HSV-1 and HCMV particles to a similar relative extent (**Figure 3A, Supplementary Table 5**).

**Figure 3:**
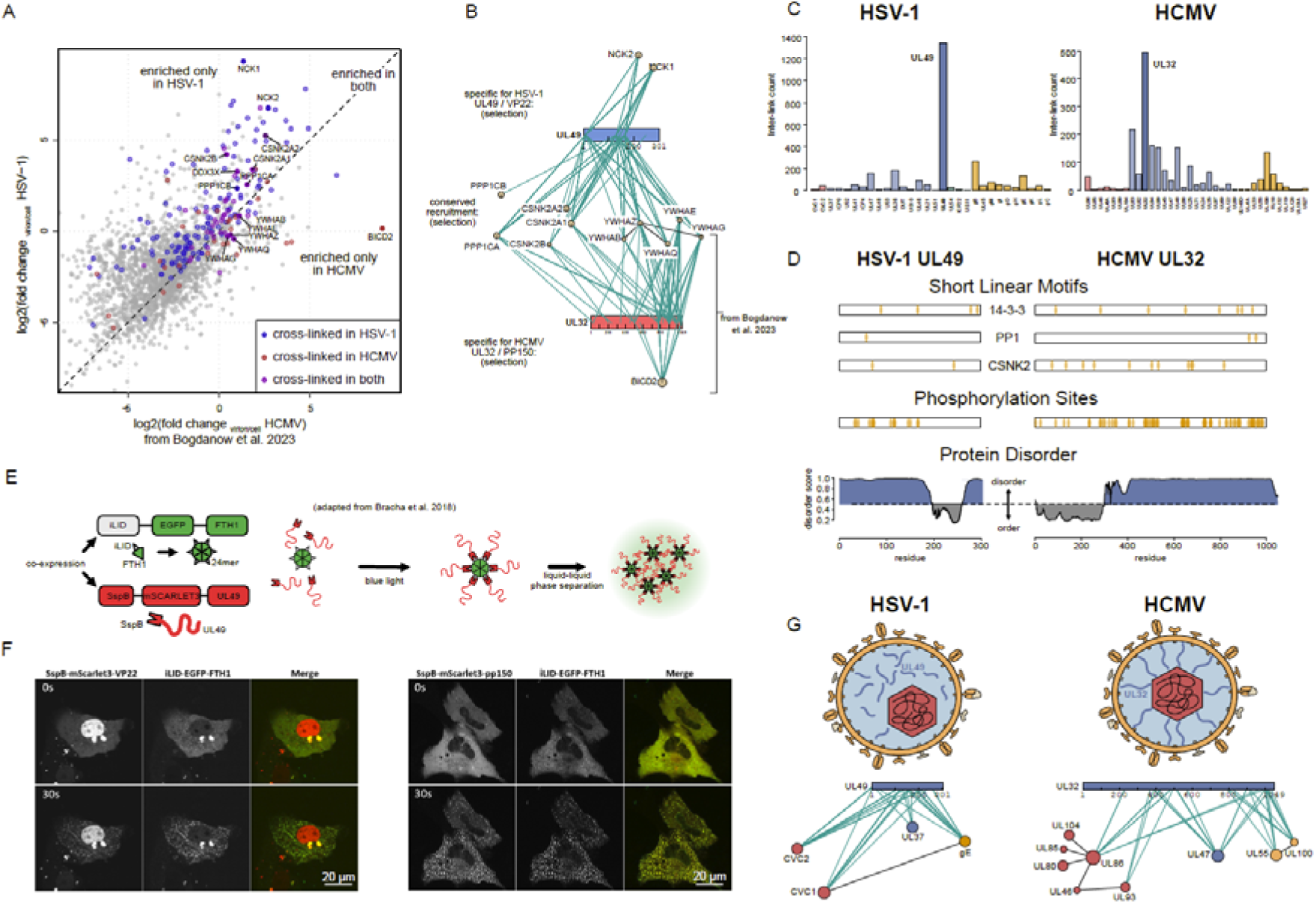
Conserved and specific tegument associations in HSV-1 and HCMV. **A:** Scatterplot comparing the relative enrichment (log2 fold change of LFQ values, virion over infected cell) of host proteins into HSV-1 and HCMV virions. Only proteins with enrichment values in both datasets are displayed. Points are highlighted in color if they were detected in the HSV-1 XL-MS dataset (blue), in the HCMV XL-MS data set (red) or in both (purple). Selected host proteins that have either similar or very different enrichment behavior are further highlighted by darker color and labeled by gene name. The HCMV virion XL-MS data are derived from Bogdanow et al. 2023 (*11*). **B:** Residue-specific cross-linking sub-network of tegument protein UL49 (HSV-1) and UL32 (CMV) with the highlighted host proteins from A. Host proteins are grouped if cross-linked only to UL49 (above), only to UL32 (below) or to both in each dataset (middle). Residue-specific representation of cross-links was done with XiNet (*67*). **C:** Number of inter-links (cross-links to other proteins) per viral protein in the dataset presented here (HSV-1, left) or from Bogdanow et al. 2023 (*11*) (CMV, right). UL49 (HSV-1) and UL32 (CMV) are highlighted. **D:** Comparison of selected shared characteristics of UL49 and UL32. Both proteins are represented as a linear bar from N- to C-terminus. Positions of similar short linear motifs (14-3-3 derived from 14-3-3-pred (*68*), PP1 derived from Bogdanow and Mühlberg et al. 2025 (*12*) for UL49 and from Bodanow et al. 2023 (*11*) for UL32, Casein kinase 2 derived from ELM) as well as phosphorylation sites (derived from Kulej et al. 2017 (*21*) for UL49 and from Bogdanow et al. 2023 (*11*) for UL32) are highlighted. Predicted local protein disorder (derived from AIUPred (*69*)) is represented as line plots. **E:** Schematic representation of workflow for in vitro testing of liquid-liquid-phase separating (LLPS) character of target protein (UL49) using the corelet system. In this system, the protein of interest is fused to SspB and mCherry-tagged. SspB’s cognate partner iLID is co-expressed fused to GFP-tagged ferritin core that oligomerizes as 24-mer. Upon light induction, iLiD is interacting with SspB and droplets become visible under fluorescence microscopy, if the protein of interest undergoes LLPS. **F:** representative images of UL49 and UL32 undergoing LLPS in the corelet system, 30 seconds after light induction of iLiD-SspB interaction. Scale bar represents 20 µm. **G:** Schematic representation of difference in cross-link pattern of UL49 and UL32 among other structural proteins (capsid, inner tegument and glycoproteins). Residue-specific representation of cross-links was done with XiNet (*67*).

Several similarly recruited host proteins are related to regulation and/or binding of phosphoproteins, including kinases (CSNK2A/B), phosphatases (PPP1CA/B), and 14-3-3 proteins (YWHAx, with x being either Q,B,Z,G,E). These proteins cross-linked to UL49 in HSV-1 and UL32 in HCMV, both of which are subfamily-unique tegument proteins of alpha- or betaherpesviruses **(Figure 3B)**. UL32 and UL49 are the most connected proteins in the network, with the majority of proteins cross-linked to either of them **(Figure 3C)**. Thus, both HSV-1 and HCMV contain subfamily-unique tegument proteins as central interaction hubs in the virion network and shared phospho-regulatory host interaction partners.

These similarities may be explained by (i) similar binding motifs in the sequence, (ii) shared biophysical or structural characteristics, or (iii) both. We have previously identified PP-1 binding motifs embedded in the disordered regions of UL32(*11*) and UL49(*12*). Moreover, both proteins contain validated or predicted short linear motifs (SLiMs) for binding to 14-3-3 proteins, feature consensus casein kinase 2 phosphorylation sites, and have been found to be phosphorylated (*11, 21*) (**Figure 3D**).

We then asked whether both proteins have common biophysical characteristics in addition to their binding motif similarities. Since UL32 is able to undergo liquid-liquid phase separation (LLPS) *in vitro* (*22*), we conducted a similar experiment with HSV-1-UL49 using the Corelet system (*23*) and compared it to HCMV-UL32 (**Figure 3E**). We observed that UL49 was able to form droplets within 30 seconds of photoactivation (**Figure 3F**), mirroring the behavior of the UL32-IDR (*22*). Both the disordered N-terminal region and the structured part of UL49 were required to undergo LLPS (**Supplementary Figure 3**). Collectively, UL32 and UL49 share similar sequence motifs for interaction with host proteins, similar network topological positions, as well as biophysical and structural characteristics, suggesting convergent tegument-organizing function of both proteins.

While these data establish HSV-1-UL49 and HCMV-UL32 as conserved at the level of network logic, there are also important differences in the structural organization inside the virion. HCMV-UL32 is nucleocapsid anchored (*24*) and spans the tegument in an N- to C-terminal radial orientation (*11*), which is reflected by the defined accumulation of cross-links to capsid proteins in the N-terminal part, glycoproteins UL55 and UL100 in the C-terminal part and the inner tegument protein UL47 to both. In contrast, HSV-1-UL49 is not nucleocapsid-associated. Accordingly, its cross-links to the corresponding HSV-1 proteins CVC1/2 (capsid), UL37 (inner tegument) and gE (envelope) are not domain-specific but spread across the entire protein (**Figure 3G**). Moreover, we detected cross-links of gE to CVC1 (and in one replicate also to CVC2, **Supplementary Table 1**).

Collectively, this suggests that UL49 does not adopt a similar tegument-spanning structural function for protein recruitment like UL32. This, as well as the CVC1/2-gE cross-links are in line with a microscopy-supported model of an asymmetrically placed HSV-1 capsid directly contacting the envelope and a tegument “cap” on the opposing side (*25–27*). Thus, HCMV-UL32 and HSV-1-UL49 are major abundant scaffold tegument proteins that share structural and biochemical characteristics but are embedded in virus-specific architectural context.

### Maturation-dependent interactome remodeling

The position of UL49 as the dominant network hub in the virion raised the broader question of how the virion network is generated. In principle, the network could reflect abundant intra-cellular associations present late in infection or, alternatively, a selection of specific interaction states from the infected cell landscape. To discriminate between these two possibilities, we compared our intra-virion PPI dataset to the previously generated infected-cell SHVIP map as a reference landscape generated using the same cell type (*12*) (**Supplementary Figure 4, Supplementary Table 6**). Overall, we observed that 117 out of 239 (49 %) interactions corresponding to ∼ 60 % of the host proteins within the particle can already be observed in intact infected cells **(Figure 4A)**. In contrast, much fewer interactions from infected cells were observed in virions (∼15%). This is expected because the intracellular network also captures processes not happening in virions, such as replication or nuclear egress.

**Figure 4:**
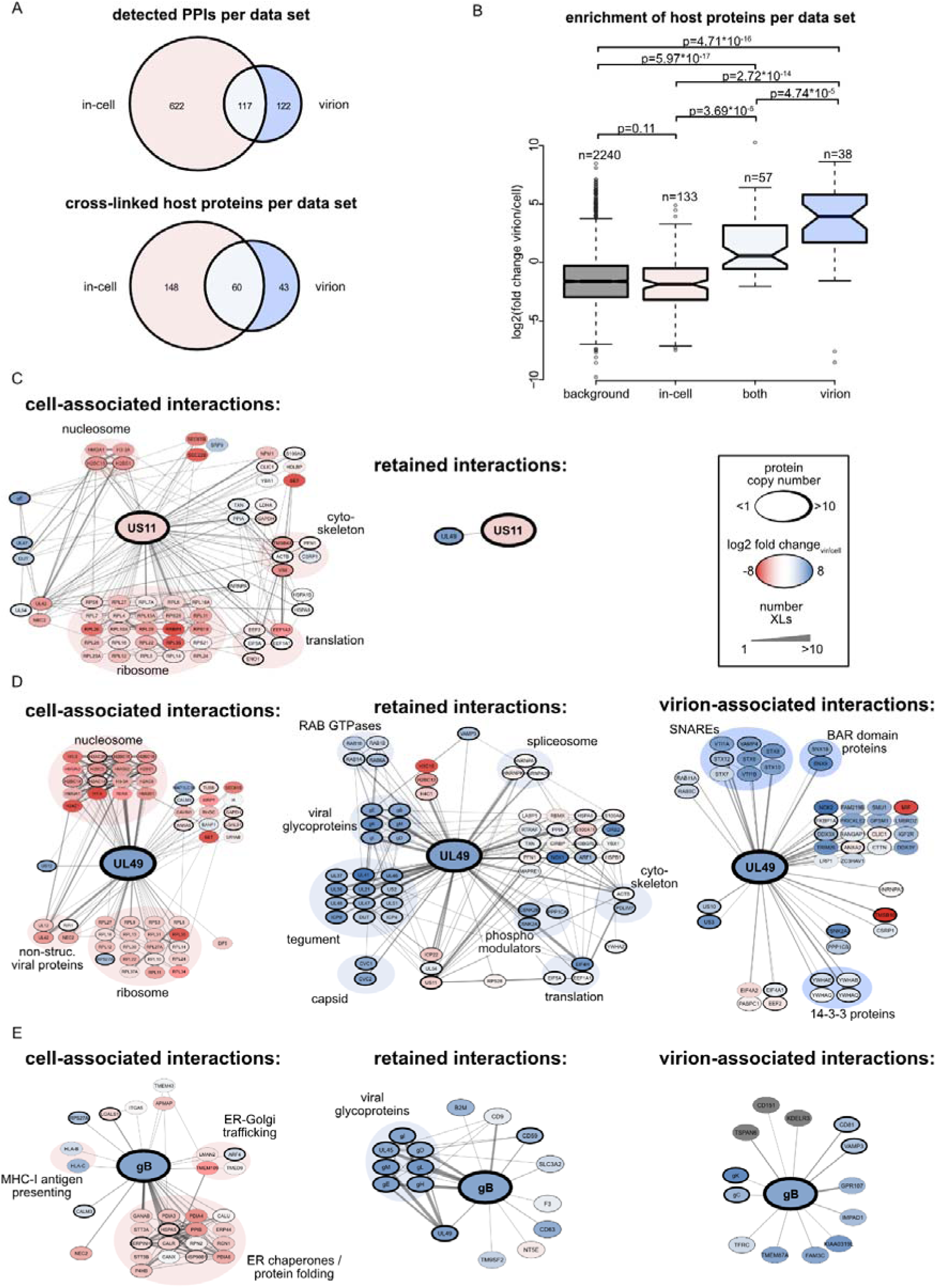
Virion maturation selectively filters the infected-cell interaction landscape. **A:** Venn diagram showing the number of PPIs (above) and of the corresponding host proteins (below) that have been detected in the XL-MS data sets of HSV-1 at a late infection stage (SHVIP, from Bogdanow and Mühlberg et al. 2025 (*12*)), in the mature virion (this work) and in both. **B:** Enrichment values (virion over cell, log2 fold change of LFQ values, see Figure 2) of the host proteins of the different subsets in A, as well as all non-cross-linked proteins detected and quantified in virions and infected cells (background). P-values showing statistical significance are shown (two-sided Wilcoxon rank sum test without multiple hypothesis correction). Boxes indicate lower and upper quartiles, horizontal line the median and whiskers extend to 1.5 times interquartile range with outliers shown. **C-E:** Subnetworks of PPIs of non-structural HSV-1 protein US11 (**C**), tegument protein UL49 (**D**) and glycoprotein gB (**E**) and among all the corresponding interactors detected exclusively at assembly stage (“lost”, left), detected in both the intracellular and the virion stage (“maintained”, middle) or exclusively detected in the mature virion (“gained”, right). Edge width scales with the number of detected cross-links. Nodes are colored according to their enrichment values (virion over cell, log2 fold change of LFQ values) from red (depleted from mature virion) over white (equal distribution) to blue (enriched with virion). For grey nodes no LFQ ratio was built due to missing data points. Thickness of node borders represents protein copy number per viral particle (see supplementary Figure 2B). Selected proteins are grouped according to protein functions or complexes.

To assess whether the interactions missing in either virion or in-cell datasets are due to technical reasons, such as limited depth of the analysis, or genuinely enriched in either cell or virion context, we correlated the virions versus cell enrichment ratios from our earlier experiments (see **Figure 2**). We observed that proteins uniquely captured in the in-cell network had approximately the same enrichment ratios as non-cross-linked host proteins **(Figure 4B)**. In contrast, host proteins captured in both, in-cell and virion datasets were significantly more enriched, and host proteins only captured in virions even exceeded these ratios. Thus, although XL-MS datasets are inherently incomplete, the enrichment behavior of these protein classes argues that many stage-specific interactions reflect genuine remodeling of the HSV-1 interactome during maturation rather than limited detection depth alone.

This remodeling process is exemplified by the non-structural viral protein US11, a known and abundant virion component (∼ 80 copies, see **Supplementary Table 2**)(*28*). In cells, US11 is cross-linked to ribosomes, confirming previous findings (*28*), as well as cytoskeletal proteins and nucleosomes, but none of these host interactions is present in virions **(Figure 4C)**. By contrast, US11 is packaged, likely due to an interaction with UL49, an interaction consistently observed in virions and in-cell datasets. This indicates that only molecules of US11 free of cellular interactors are packaged into the virion. A similar pruning of interactions is also observed for other non-structural viral proteins such as UL54 or ICP22, which are incorporated into virions while depleted from nucleus-residing host and viral interactors **(Supplementary Figure 4)**. Thus, few non-structural viral proteins get incorporated into newly formed virions but lose almost all their intra-cellular interactions.

We next asked how the interaction-states of structural proteins are changed during maturation and specifically investigated gB and UL49. In cells, UL49 had many interactions across cytosolic and nuclear compartments, including nucleosomes and ribosomes. These interactions get lost in virions **(Figure 4D)**. In contrast, some interactions to UL49 are gained in the virion. These involve membrane remodeling factors such as v- and t-SNAREs (STX7, STX12, VTI1A, VTI1B, STX6, STX8, STX10, VAMP4), which function in vesicle fusion at the Golgi or endosome compartments (*29*) and PX-BAR-domain-proteins (SNX9 and SNX18), known to shape membrane curvature (*30, 31*). A core UL49-interactome involving other tegument proteins, glycoproteins, RAB GTPases, cytoskeletal and phospho-regulatory proteins is found in both virions and infected cells.

A similarly extensive remodeling is observed for the altogether smaller gB interactome. During maturation, gB lost intracellular interactors related to processing of glycoproteins, such as ER-chaperones, but gained proteins related to vesicle fusion **(Figure 4E)**. Taken together, virion maturation is accompanied by extensive network remodeling, involving loss of the majority of intracellular interactions, maintenance of a core assembly interactome, and recruitment of potentially infection-relevant proteins such as membrane remodeling factors.

### Selective incorporation of CD59 links virion maturation to complement evasion

Among the gB interactions present in both cells and virions is CD59, one of the most enriched host factors within the virion (**Figure 2**). This protein was enriched in HSV-1 virions but undetected in HCMV virion and cell samples (**Figure 3A**). CD59 is an inhibitor of complement-mediated lysis, a mechanism of the innate immune system which leads to the clearance of outer membrane-containing pathogens after being bound by an antibody (*32, 33*). CD59 inhibits the last step in the complement cascade: the formation of the membrane attack complex (MAC) (*34*). In support of this function, cross-links detected between CD59 and the extracellular domains of viral membrane proteins gB and UL45 indicate that this host protein is located on the outside of the viral envelope **(Figure 5A, B)**. To test whether CD59 on the virion surface might have a protective function for HSV-1, we blocked CD59 function using a specific antibody (BRIC229) (*35*) in the presence of complement-containing serum **(Figure 5C)**. We treated HSV-1 virions prepared from infected cell culture supernatant with HSV-1 seronegative human complement-containing serum in the presence of BRIC229 and complement-activating, non-neutralizing gH-antibody BBH-1. We found that specific blocking of CD59 via BRIC229 renders HSV-1 particles sensitive to complement treatment which manifested in a significant reduction of viral infectivity **(Figure 5D, Supplementary Table 7)**. Prior complement inactivation by heating or omitting the activating antibody reversed this effect. Thus, HSV-1 interacts with CD59 during intracellular stages - an interaction that is maintained during assembly and results in recruitment of CD59 to the virion particle surface, which protects HSV-1 from complement-mediated inactivation.

**Figure 5:**
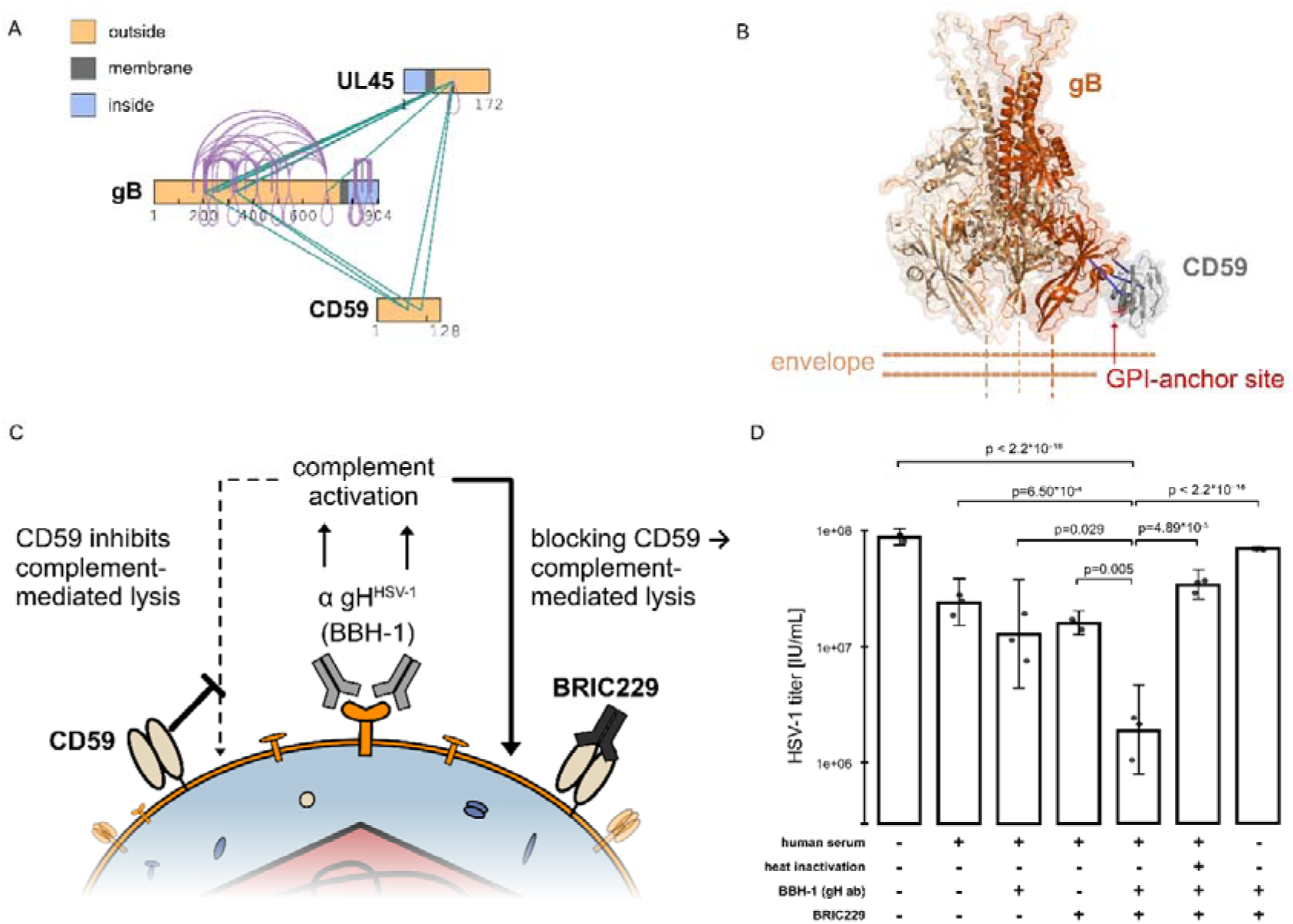
The host protein CD59 protects HSV-1 against complement-mediated lysis. **A:** CD59 was found cross-linked to virion-surface domains of viral envelope proteins gB and UL45. Residue-specific representation of cross-links was done with XiNet (*67*). **B:** Co-prediction of CD59 (grey) and homotrimeric gB (different shades of orange) with AF3x (*70*) using a subset of intra- and inter-links shows the contact site between the host factor and the viral glycoprotein in pre-fusion conformation close to the viral membrane. gB transmembrane domains are not included in the prediction and are emphasized by dashed lines. GPI-anchor attachment site of CD59 is shown in red. Three inter-links between CD59 and gB used to guide the prediction are shown in blue. **C:** Schematic workflow for testing the function of CD59 on the viral envelope in protecting HSV-1 against complement-mediated lysis. After binding of a gH-specific antibody (BBH-1) and treatment with complement-containing serum the complement cascade would lead to lysis of HSV-1, which is inhibited by CD59. When CD59 is blocked by a specific antibody (BRIC229) lysis is facilitated. **D:** Testing the function of CD59 on the viral envelope. Equal parts of extracellular infectious HSV-1 virions were treated with (HSV-1 seronegative) complement containing human serum and different combinations of complement activating, non-neutralizing gH antibody (BBH-1), CD59 inhibiting antibody (BRIC229), or with prior heat inactivation of the serum. Effect of treatment on HSV-1 was tested by titrating the samples over uninfected fibroblast cells and quantifying viral titers by measuring ICP4 expression after 12 h by FACS (FFU/mL = focus forming units per milliliter). All samples were made in biological triplicates, which are shown as dots with bar height indicating the mean and whiskers standard error. Statistical significance was tested by one-way ANOVA and Dunnett’s multiple-comparisons adjustment.

## Discussion

How complex viruses transform intracellular interaction networks into an infectious extracellular particle has remained poorly defined. By integrating virion XL-MS, quantitative proteomics, and infected-cell interactomics, we show that HSV-1 particle maturation is a highly selective process that operates not only at the level of protein incorporation but also at the level of molecular associations. Mature virions therefore do not represent a passive snapshot of the late stage infected cell. Rather they constitute a remodeled subset of the intracellular HSV-1 interaction landscape that is enriched for associations compatible with particle architecture, envelopment, and extracellular function.

Non-structural proteins, such as US11, illustrate this principle. US11 is incorporated into virions, yet many of its dominant intracellular associations are lost during maturation. For example, the strong association with 60S ribosomal subunits (*28*) is absent from mature particles, indicating that virion incorporation is selective for specific molecular states rather than for proteins per se. These observations suggest that herpesvirus maturation acts as a molecular filter that enriches interaction states compatible with virion function while excluding associations linked to intracellular biosynthetic activities.

This remodeling extends to structural proteins, such as the abundant tegument protein UL49 - the most connected node. We find that UL49 PPIs are extensively remodeled during maturation: chromatin- (*36, 37*) and ribosome-associated (*38*) intracellular interactions are depleted, whereas mature virions retain a compact UL49-centred network linking viral structural proteins to host factors involved in membrane traffic and remodeling, including SNAREs and PX-BAR domain proteins. The latter may add to mechanistic models of HSV-1 envelopment, which are currently centered on associations among viral proteins (*39*) and involvement of ESCRT proteins (*40, 41*).

Previous studies showed that UL49 contributes to incorporation of tegument and glycoproteins (*42, 43*), with cell-type and system-dependent additional roles in growth (*42*), spread (*43*), host shutoff (*44*) and plaque-phenotypes (*45*). Our work extends this view by assigning UL49 a multivalent physical hub role, positioning it as the most dominant interface between the viral and host proteome. This may be relevant for host-cell imprinting of newly produced viruses by recruiting cell-type-specific host factors, influencing downstream infectivity phenotypes (*46*).

Comparison with HCMV (*11*) reveals that this organizational principle extends across divergent herpesvirus lineages. HSV-1 UL49 and HCMV UL32 are non-orthologous, lineage-specific tegument proteins, but both occupy highly connected positions within their respective virion networks. Their shared dependence on phosphoregulatory host factors, short linear interaction motifs and LLPS (*22*) suggests convergence at the level of tegument network organization and assembly mechanism. This is reminiscent of other cases of herpesviral functional convergence, such as the independent evolution of immunoevasins targeting the antigen-presentation pathway (*47*).

The property to tune LLPS through phosphorylation and de-phosphorylation enables HCMV-UL32 to regulate assembly processes (*22*), such as by recruiting membrane-associated viral proteins (*22*). In HSV-1, UL49 could have a similar role through its interactions with viral glycoproteins (*48*) and host-derived membrane-remodeling factors. Notably, these similarities in network organization persist despite major differences in molecular implementation, including the stable capsid association of UL32 (*19*), which is absent for UL49.

We also observe extensive remodeling at the outside of the membrane, For example, interactions of viral glycoproteins with host proteins related to folding and processing of glycoproteins in the ER become depleted from virions, consistent with a loss of immature biosynthetic states during glycoprotein maturation (*49, 50*). Instead, we observe abundant and selective incorporation of complement inhibitor CD59 on the virion surface and propose this as an immune-evasion mechanism that protects from complement-mediated loss of infectivity, as described for several unrelated enveloped viruses (*51–55*). HSV-1 is known to antagonize the complement cascade by binding of antibody Fc domain by gE (*6, 56*) and by inhibition of the cleavage cascade by binding the C3b fragment via gC (*57*). The mature HSV-1 particle therefore carries a layered complement-evasion system, combining viral Fc/C3b interference with host-derived MAC inhibition, indicating that maturation also refines for extracellular immune escape. Taken together, our data define herpesvirus maturation as a selective remodeling process that couples tegument organization, convergent network architecture and complement evasion within the mature infectious virion.

## Methods

### Cells and viruses

Human embryonic lung fibroblasts (HELFs, Fi301, obtained from the Institute of Virology, Charité, Berlin, Germany) were maintained in a monolayer culture as described before (*58*). To reconstitute cloned HSV-1 strain 17^+^, cells were nucleofected with the bacterial artificial chromosome containing the viral genome (*59*) by electroporation. After cytopathic effect was pronounced for all cells (about 24 hours after infection), cell culture supernatant was harvested and the amount of infectious particles was determined by immunotitration as described below.

### Virion harvest, cross-linking and purification

HSV-1 was harvested from infected fibroblast cell culture supernatant about 24 hours post infection. Cellular debris was removed by centrifuging the fluid at 1500 g. Subsequently, virions were pelleted by centrifuging the supernatant in a SW-28 rotor (Beckman) at 25,000 rpm for 1 h at 10°C. After the pellet was washed once, virions were resuspended in PBS and cross-linked by addition of DSSO cross-linker dissolved in DMSO to reach a concentration of 3 mM. Cross-linking was carried out for 30 min at room temperature while shaking after which the reaction was quenched by addition of 50 mM Tris-HCl. Cross-linked virions were loaded on a glycerol tartrate density gradient (*60*) and centrifuged in a SW-40 rotor for 1 h at 10°C with brakes set at lowest deceleration. Virion band was taken off using a 1.2 mm syringe and the resulting sample was washed twice with PBS to remove the gradient material, pelleting virions first using a SW-60ti rotor at 30,000 rpm and second using a TLS-55 rotor at 35,000 rpm, 1h and 10°C respectively. Resulting samples were stored as pellets in −80 °C until further sample preparation.

### Sample preparation for bottom-up proteomics

For preparation of HSV-1 samples for bottom-up proteomics virion samples were prepared as described above with the exception of skipping the cross-linking before gradient centrifugation. During the washing steps after purification, particle concentrations of all samples were adjusted to one another by measuring OD600. For comparison with intracellular protein levels an aliquot of the infected cells the supernatant was taken from was also harvested, washed with PBS two times and stored as pellets at −80°C. All bottom-up samples destined for label-free quantification were prepared in four or three biological replicates respectively.

### Protein extraction and digest from virions and cells for XL-MS or bottom-up proteomics

Samples were resuspended in PBS and glycoproteins were deglycosylated using a deglycosylation kit (Protein Deglycosylation Mix II, New England Biolabs, denaturating conditions). After this, lysis buffer containing 8 M Urea in 50 mM triethylammonium bicarbonate (Teab), supplemented with cOmplete protease inhibitor (Roche) was added to the sample. Genomic DNA was removed by addition of Benzonase. For cross-linked virion samples lysis was further facilitated by sonication with a Sonopuls HD 2070 Sonicator equipped with a MS73 Sonotrode (Bandelin) with 0.5 s intervals of sonication at 50% intensity and pause for 30 s while keeping the samples on ice. Bottom-up samples were sonicated using a Bioruptor Pico with 10 cycles alternating between 30 s sonication and 30 s pause while cooling the sample at 4°C. After that, proteins were extracted by methanol/chloroform precipitation (*61*) and finally resuspended in digestion buffer containing 1% (w/v) sodium deoxycholate, 5 mM dithiothreitol and 40 mM chloroacetamide in 50 mM Teab. Proteins were digested by addition of trypsin at an estimated enzyme to protein ratio of 1:100. Digestion was carried out overnight at 37°C while shaking. Finally, digestion was stopped by acidification of the sample with 1% (v/v) formic acid (FA) and peptides were desalted using Sep-Pak C8 cartridges (Waters) in case of cross-linked samples or using c18 stage-tips (*62*) in case of bottom-up samples. Samples were dried in speed-vacuum and stored at −20 °C until further preparation or LC-MS analysis.

### Off-line HPLC fractionation of cross-linked peptides

Cross-linked peptides were separated from linear peptides by strong cation exchange (SCX) chromatography. Therefore, approximately 100 µg of cross-linked peptides were resuspended in 0.1% (v/v) FA and loaded on a PolySULFOETHYL A column (PolyLC) connected to an Agilent 1260 Infinity II system. The peptides were separated by a 95 min linear gradient going from buffer A (0.1% v/v trifluoroacetic acid, TFA in 20% v/v acetonitrile, ACN) to buffer B (0.5 M NaCl and 0.1% v/v TFA in 20% v/v ACN) using the following increments: 0 min 0% buffer B, 0.01 min 2% buffer B, 8.01 min 3 % buffer B, 14 min 8% buffer B, 28 min 20% buffer B, 48 min 40% buffer B, 68 min 90% buffer B, 74 min 90% buffer B, 75 min 0% buffer B, 95 min 0 % buffer B, flow rate 0.18 mL/min. Fractions were collected in 60 s intervals, dried and desalted using C8 stage-tips. Desalted samples were dried and stored at −20 °C until measurement.

### LC-MS measurement of cross-linked samples

Cross-linked peptides were analyzed with an Orbitrap Fusion Lumos Tribrid mass spectrometer (Thermo Fisher Scientific) equipped with a FAIMS Pro Duo interface and controlled by instrument control software version 4.0. Approximately 1 µg of peptides was loaded onto a 50 cm in-house packed analytical reverse-phase column (Poroshell 120 EC-C18, 2.7 µm, Agilent Technologies). Separation was achieved with 180 min linear gradients from 0.1% FA in water (buffer A) to 0.1% FA in 80% ACN (buffer B) at a flow rate of 250 nL/min. Data were acquired using a stepped-HCD-MS2 method with alternating FAIMS compensation voltages between −50, −60, and −75 V. MS1 scans were recorded in the Orbitrap at a resolution of 120,000 with the following parameters: scan range 375–1600 m/z, AGC target set to “Standard,” maximum injection time of 50 ms, precursor charge states +4 to +8, isolation window of 1.6 m/z, dynamic exclusion of 60 s, and a cycle time of 2 s. Fragmentation was performed using stepped HCD energies of 21%, 27%, and 33%. MS2 scans were acquired in the Orbitrap at a resolution of 60,000 with mass range set to “Normal” and scan range set to “Auto.” The maximum injection time was 118 ms and the AGC target was set to 200%.

### LC-MS measurement of bottom-up samples

Bottom-up samples were analyzed with an Orbitrap Exploris 480 mass spectrometer (Thermo Fisher Scientific) coupled online to a Vanquish Neo UHPLC system (Thermo Fisher Scientific) and controlled by instrument software version 4.2. Peptides were loaded onto a 50 cm in-house packed analytical reverse-phase column (Poroshell 120 EC-C18, 2.7 µm, Agilent Technologies) and separated by a 180 min linear gradient from 0.1% FA in water (buffer A) to 0.1% FA in 80% ACN (buffer B) at a flow rate of 250 nL/min. MS1 scans were acquired in the Orbitrap at a resolution of 120,000 with a cycle time of 2 s and a dynamic exclusion of 40 s. The precursor intensity threshold was set to 1E+4, with the maximum injection time set to “Auto” and an AGC target of 300%. Precursors with charge states +2 to +4 were isolated using a 1.6 m/z window and fragmented with HCD at 30%. MS2 scans were acquired in the Orbitrap at a resolution of 15,000 with the AGC target set to “Standard.”

### XL-MS data analysis

Cross-linked peptides were searched using Scout v1.4.1 (*63*) using standard parameter settings for DSSO cross-linker. This includes: cross-linker mass 158.00376533 Da, light fragment 54.01056468 Da, heavy fragment 85.98263585 Da, cross-linked residues K,S,T and Y as well as the protein N-terminus. Additional search parameters include carbamidomethylation of cysteines (+57.02146 Da) as a static modification, oxidation of methionine (+15.9949 Da) and acetylation of protein N-terminus (+42.0106 Da) as variable modifications, 7 amino acids minimum peptide length, 3 maximum missed cleavages, 500 Da minimum peptide mass, 6000 Da maximum peptide mass, 10 ppm MS1 mass error, 20 ppm MS2 mass error and trypsin as the digestion enzyme. Results were filtered to 1% FDR on CSM, residue-pair and PPI-level separating FDR calculation between intra-and inter-cross-links. In case of peptides mapped to two partially homologous proteins, the protein with more overall identified cross-links was selected as the linked protein. If the two partially homologous proteins had the same frequency of detected cross-links, the protein with the higher iBAQ in the bottom-up analysis was selected.

### Bottom-up proteomics and label-free quantification data analysis

Raw files were searched with MaxQuant version 1.6.2.6a (*64*) against the Uniprot human reference proteome (TaxonID 9606, downloaded 2020) and the Uniprot reference proteome of Human herpesvirus 1 strain 17 (Human herpes simplex virus 1, Taxon ID 10299, downloaded 2020). Standard parameters were used for the search and match between runs as well as second peptides were enabled. Trypsin was selected as the digesting enzyme, minimum peptide length was set to 7 amino acids and maximum of 2 missed cleavages were allowed. iBAQ quantification was enabled. Results were filtered to 1% false discovery rate (FDR) on PSM and Protein level using reversed decoy sequences. Protein intensities, iBAQ values or LFQ-values were analyzed from proteinGroups.txt file using R. For label-free quantification analysis, missing values were imputed according to (*65*) with random numbers sampled from a normal distribution shifted 1.8 standard deviations downward and with a distribution width of 0.3 relative to each sample. Fold changes were calculated from these values and log2-transformed. p-values were calculated by students t-test in R. For comparative analysis of HSV-1 and HCMV enrichment levels, additional raw files containing HCMV virion versus cell experiments were used from Bogdanow et al. 2023 (*11*) and all raw files were searched in a combined MaxQuant run. LFQ intensities were furthermore median-normalized prior to building fold changes. iBAQ values from virion samples were used for absolute copy number estimations using known copy numbers of capsid proteins as an internal standard: UL19 955 copies, scp 900 copies, UL18 640 copies, UL38 320 copies, UL36 120 copies, CVC2 120 copies, CVC1 60 copies and UL6 12 copies per capsid and virion. Slopes and intercepts of linear regressions between iBAQ values and protein copy numbers, plotted on double-logarithmic scales, were estimated in R and subsequently used to calculate copy numbers for the remaining proteins from their iBAQ values according to the following equation:

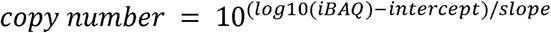

### CD59 inhibition and treatment of virions with human complement

Human complement containing blood serum was prepared from freshly drawn blood from a HSV-1 seronegative tested author of this manuscript. The blood was incubated in a glass tube at room temperature for 30 minutes to allow for blood clotting to take place. The clot was removed by centrifugation at 1000 g for 10 minutes, followed by another centrifugation step with the supernatant at 12,000 g for 10 minutes. The resulting blood serum was aliquotted, snap-frozen in liquid nitrogen and stored at −80 °C. Serum was thawed immediately before use at 34 °C or, if needed, heat-inactivated by incubation at 56 °C for 30 minutes.

For the treatment of virions, approximately 1E+8 infectious units of HSV-1 strain 17 per sample were prepared by ultracentrifugating infectious cell culture supernatant at 25,000 rpm and 10 °C for 1 h using a SW40 rotor. The pellets were resuspended in PBS supplemented with 0.5 mM MgCl2 and 0.15 mM CaCl2. For blocking of CD59, virion samples were pre-treated with 20 µg/mL BRIC229 (International Blood Group Reference Laboratory) or the corresponding volume of 50% glycerol as control and incubated for 30 min at 34 °C while shaking. After this, different combinations of 5% (v/v) human serum with or without previous heat inactivation and/or 5 µg/mL BBH-1 antibody (Abcam) was added to the samples, followed by incubation at 34 °C for 1 h while shaking. Infectivity of the samples was tested immediately afterwards by immunotitration.

### Immunotitration by Flow Cytometry

Virus aliquots were tested for infectivity on HELFs in at least three five-fold serial dilutions. Cells were harvested at ∼ 10 h post infection by trypsinisation, counted, washed with PBS and permeabilized in 75% (v/v) pre-cooled ethanol at 4 °C for at least 15 minutes. Cells were washed with 1% (w/v) BSA in PBS and subsequently stained for ICP4 expression using a 1:200 diluted mouse anti-ICP4 monoclonal antibody (Santa Cruz Biotechnology) followed by a 1:1000 diluted Alexa-647-coupled goat anti-mouse IgG polyclonal antibody (Invitrogen). The fraction of ICP4-positive cells was determined using a FACSCanto II flow cytometer, controlled with FACSDiva software (BD Biosciences).

### Corelet-assay for testing phase separation of UL49

Corelet plasmids were generated by InFusion cloning and sequence-verified. HSV-1 UL49 sequences were amplified from the HSV-1 HA-UL49 BAC and fused with a PCR-amplified SspB-mScarlet3 fragment in a pcDNA3.1 backbone. Construction of the pcDNA3-iLID-EGFP-FTH1 construct, as well as the SspB-mScarlet3-UL32 construct used for comparison was described previously (*22*). For transfection, VeroB4 cells were seeded in ibidi μ-Slide 8-well chamber slides (ibidi, Cat. No. 80806) to reach ∼70% confluency and transfected with 360 ng plasmid DNA per well using Lipofectamine 3000. Imaging was performed 24 h after transfection. Live-cell time-lapse fluorescence imaging was conducted on a Nikon Eclipse Ti2 spinning-disk confocal microscope equipped with a Yokogawa CSU-W1 spinning disk unit, an Andor iXon Ultra DU-888U3 EMCCD camera and a SR Apo TIRF AC 100×/1.49 NA oil-immersion objective under physiological conditions (37°C, 5% CO_2_). Corelet condensate formation was induced by 488 nm illumination, which simultaneously excited iLID-EGFP-FTH1 and triggered the iLID–SspB interaction, eliminating the need for a separate activation step. To obtain a pre-activation reference, the mScarlet3 channel was imaged before EGFP acquisition in the first frame, while all subsequent frames captured post-activation dynamics.

## Data Availability

The mass spectrometry proteomics data have been deposited to the ProteomeXchange Consortium via the PRIDE (*66*) partner repository with the dataset identifier PXD082235.

---

**Supplementary Figure 1:**
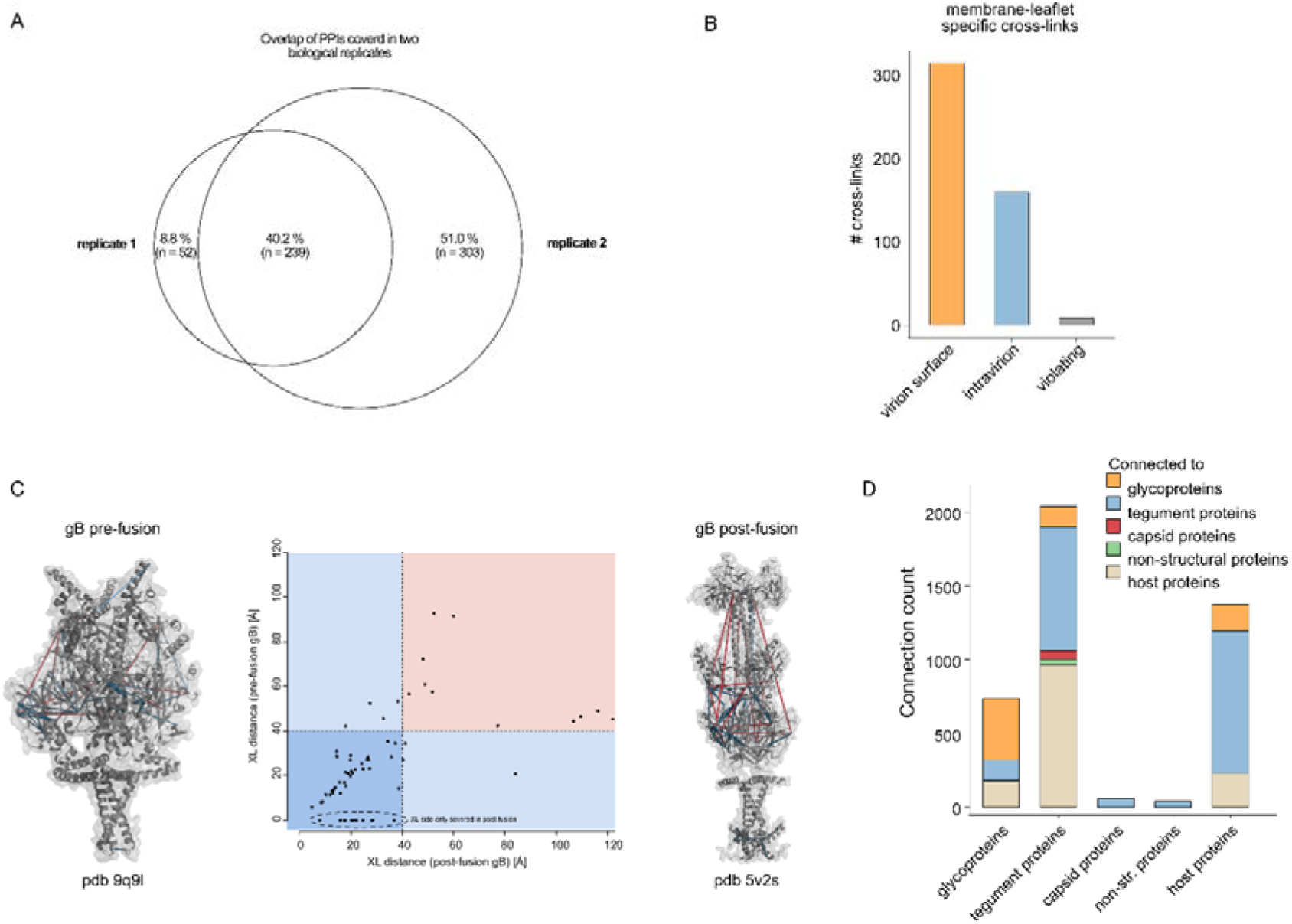
Assessment of reproducibility and agreement of XL-MS data with previous data. **A:** Reproducibility of detected PPIs in two biological replicates following the XL-MS workflow from Figure 1A. **B:** Number of cross-links (intra- and inter-links) of transmembrane proteins sorted by membrane-leaflet (virion surface, orange and intravirion, blue). Cross-links connecting protein domains from different leaflets violate the known topology of transmembrane proteins and are shown in grey. **C**: Scatterplot of cross-links within glycoprotein B plotted on the post-fusion (x-axis, PDB: 5v2s (*17*)) or the pre-fusion structure (y-axis, PDB: 9q9l (*16*)) of the homotrimeric protein. The maximum calpha-calpha distance of connected residues (40 Angström) in each structure is indicated by dashed lines. The corresponding protein structures are shown left and right in grey and cross-links are shown as lines colored according to fulfillment of the distance criteria (blue satisfied, red violated intra-link in the corresponding structure). A subset of intra-links is only plottable and satisfied in the post-fusion structure. **D:** Stacked barplot showing the number of detected cross-links connecting proteins of different classes (glycoproteins, tegument, capsid, non-structural proteins and host proteins) to the corresponding other protein classes.

**Supplementary Figure 2:**
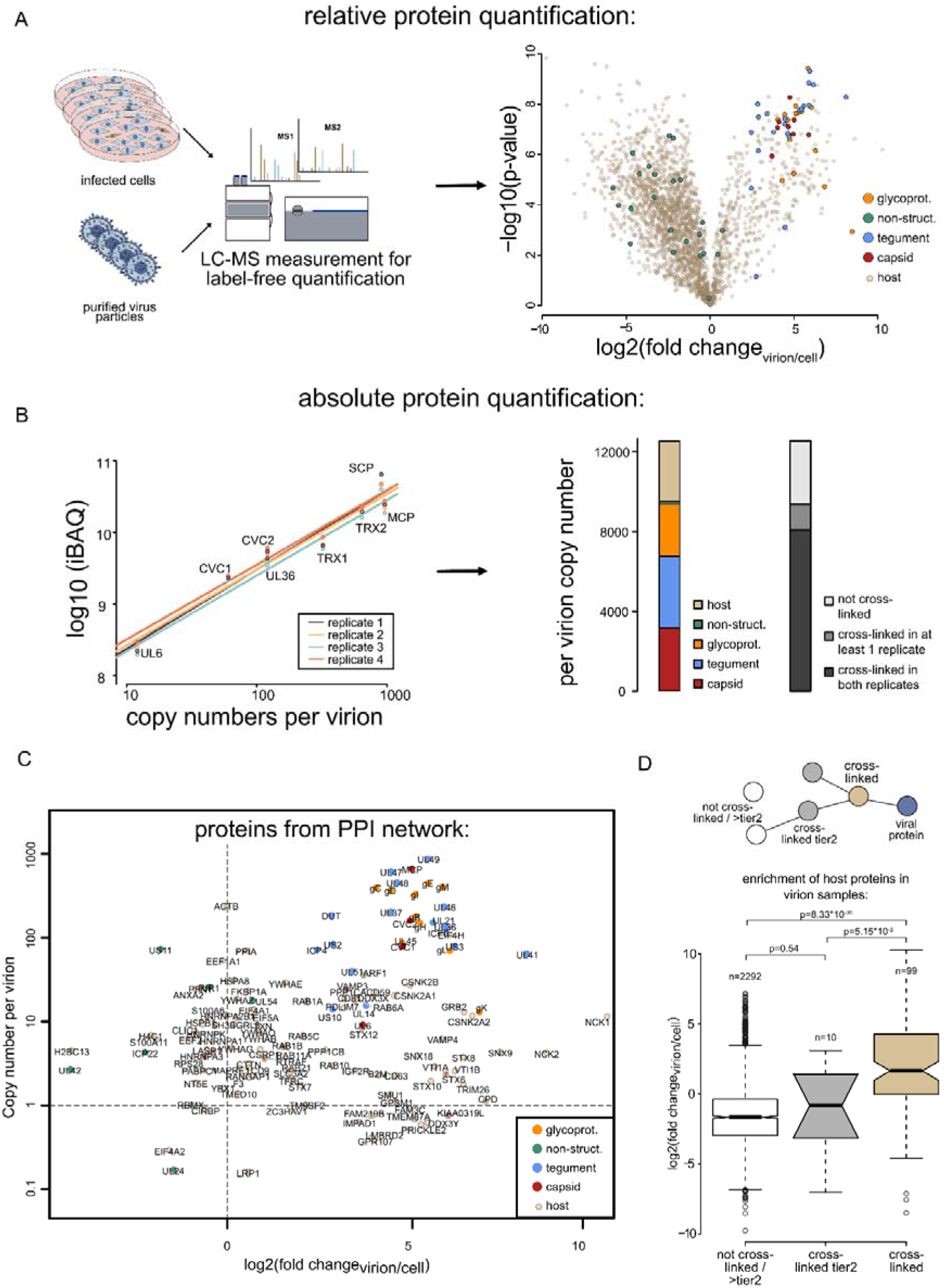
Quantifying the virion-wide proteome. **A:** Relative quantification of virion-incorporated proteins. Samples derived from purified virions or infected cells taken in biological quadruplicates are measured using label-free quantification proteomics. Results are shown in the volcano plot on the right comparing protein LFQ values showing enrichment in virions over cells (log2-transformed) with p-values representing statistical significance (students t-test, log10-transformed). Color coding according to Figure 1B. **B:** Absolute quantification of virion-incorporated proteins. Quadruplicate purified virion samples were measured as shown in A. Protein iBAQ-values (intensity based absolute quantification) of capsid proteins alongside their known capsid stoichiometries are used to make a linear regression, which derives per-virion copy numbers of all incorporated proteins based on their intensities. The sum of absolute copy numbers of all proteins per protein class from Figure 1B is shown as a stacked barplot on the right. Furthermore, the amount of proteins covered in the XL-MS data in both or at least one replicate is shown. **C:** Scatterplot comparing virion over cell enrichment (log2 fold change from LFQ values) over the absolute per-virion copy number (on a decimal logarithmic scale) of proteins from the PPI-network as shown in Figure 2. All proteins are labelled with their gene name. **D:** Boxplot comparing the enrichment values (log2 fold changes of LFQ values, virion over cell) of all detected host proteins separated into proteins directly linked to a viral protein in the PPI map, linked to a host protein linked to a viral protein (tier2) or linked over higher tiers or not at all to viral proteins. P-values showing statistical significance are shown (two-sided Wilcoxon rank sum test without multiple hypothesis correction). Boxes indicate lower and upper quartiles, horizontal line the median and whiskers extend to 1.5 times interquartile range with outliers shown.

**Supplementary Figure 3:**
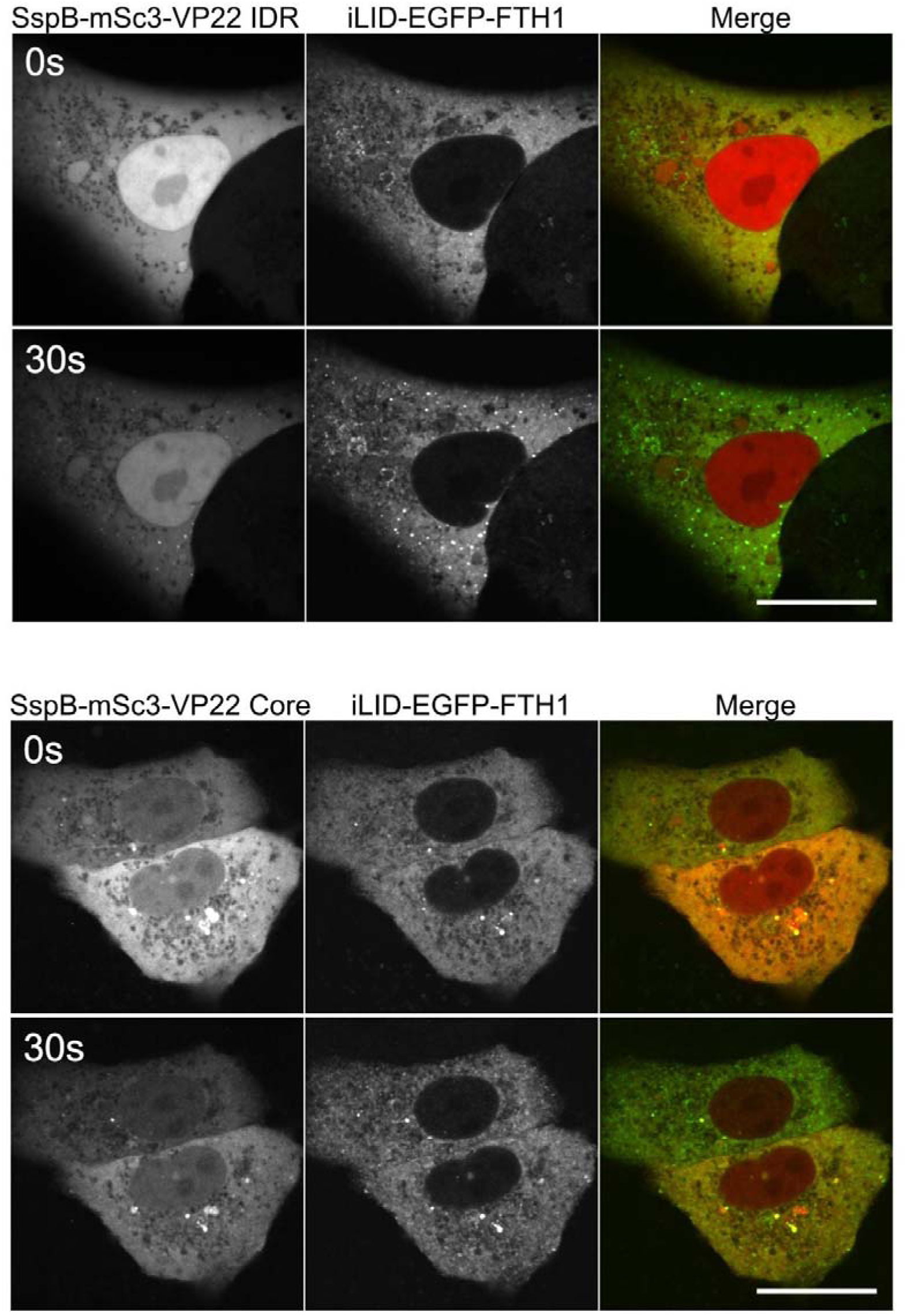
UL49 IDR domain and core domain alone are not sufficient for droplet formation. Testing of UL49 IDR domain (residues 1-173, above) and core domain (residues 173-301, below) to undergo LLPS in the corelet system. Representative pictures 30 seconds after light induction of iLiD-SspB interaction are shown. Scale bar represents 20 µm.

**Supplementary Figure 4:**
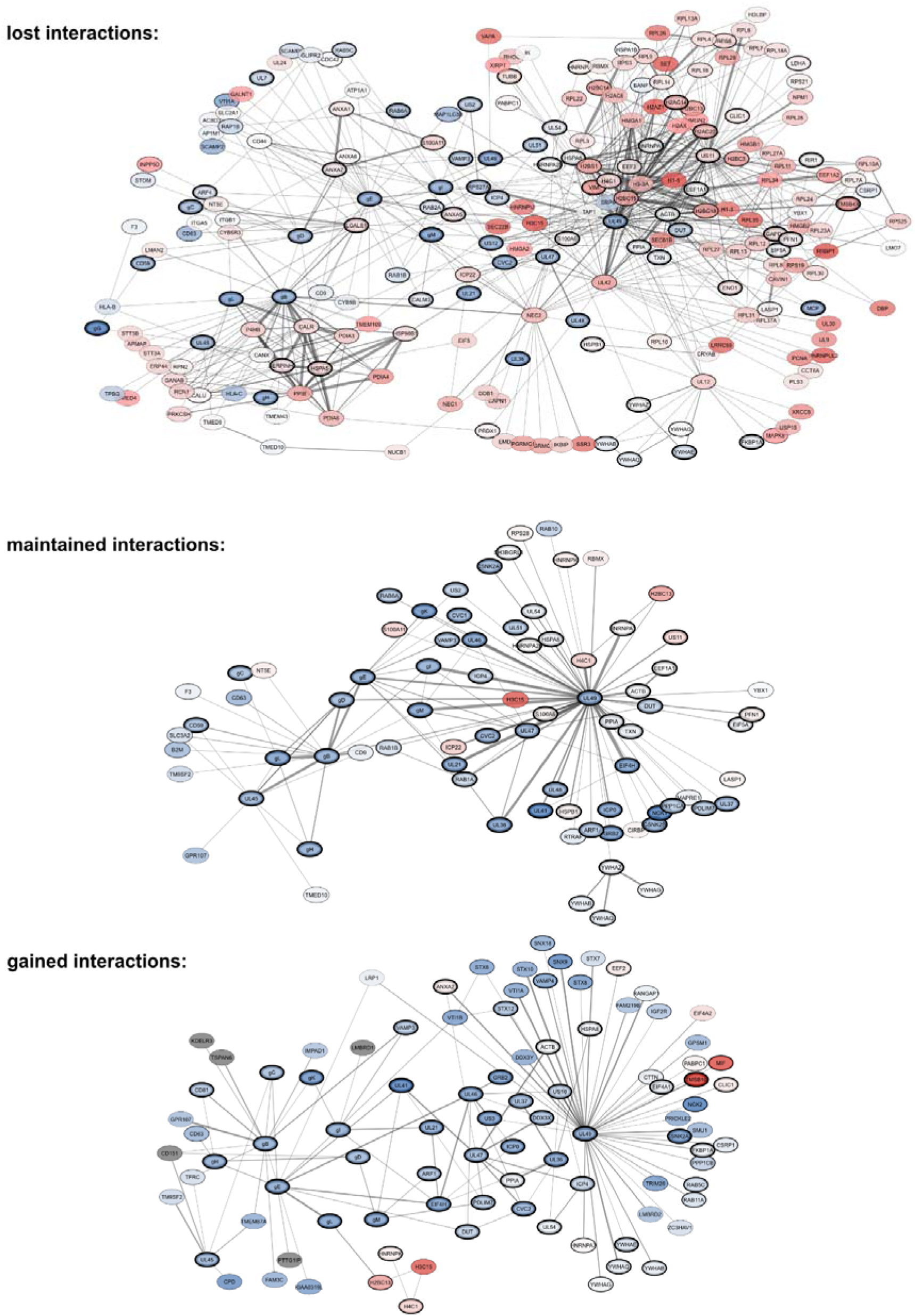
Refinement of HSV-1 protein interactome from assembly to the mature virion. Subnetworks of all PPIs among HSV-1 proteins and their direct host interactors detected exclusively at assembly stage (“lost”, above), detected in both the intracellular and the virion stage (“maintained”, middle) or exclusively detected in the mature virion (“gained”, bottom). Network representation including fold changes, copy numbers and cross-link numbers according to Figure 4 C-E.

## Author Contributions

Conceptualization: L.M. and B.B. Methodology: L.M., Y.J., J.R., I.G., J.B.B, L.W. and B.B. Investigation: L.M., Y.J., I.G., J.R., and B.B. Visualization: L.M, and Y.J. Funding acquisition: F.L., J.B.B and B.B. Formal analysis: L.M., Y.J., J.R. and B.B. Data curation: L.M. Project administration: L.M., L.W., F.L. and B.B. Supervision: J.B.B., L.W., F.L. and B.B. Resources: L.W. and F.L. Writing—original draft: L.M. and B.B. Writing—review and editing: L.M., L.W., F.L. and B.B.

## Acknowledgements

The authors thank Beate Sodeik (MHH, Hannover) for providing the seed virus and together with Florian Full (Freiburg University) for valuable comments on the manuscript.

